# Loss of *cped1* does not affect bone and lean tissue in zebrafish

**DOI:** 10.1101/2024.07.10.601974

**Authors:** Kurtis Alvarado, W. Joyce Tang, Claire J. Watson, Ali R. Ahmed, Arianna Ericka Gomez, Rajashekar Donaka, Chris Amemiya, David Karasik, Yi-Hsiang Hsu, Ronald Young Kwon

**Author notes:** Corresponding author:, Address: University of Washington, South Lake Union Campus, 850 Republican Street, Seattle, WA 98109. These authors contributed equally to this work (Kurtis Alvarado, W. Joyce Tang).

## Abstract

Human genetic studies have nominated Cadherin-like and PC-esterase Domain-containing 1 (*CPED1*) as a candidate target gene mediating bone mineral density (BMD) and fracture risk heritability. Recent efforts to define the role of *CPED1* in bone in mouse and human models have revealed complex alternative splicing and inconsistent results arising from gene targeting, making its function in bone difficult to interpret. To better understand the role of *CPED1* in adult bone mass and morphology, we conducted a comprehensive genetic and phenotypic analysis of *cped1* in zebrafish, an emerging model for bone and mineral research. We analyzed two different *cped1* mutant lines and performed deep phenotyping to characterize more than 200 measures of adult vertebral, craniofacial, and lean tissue morphology. We also examined alternative splicing of zebrafish *cped1* and gene expression in various cell/tissue types. Our studies fail to support an essential role of *cped1* in adult zebrafish bone. Specifically, homozygous mutants for both *cped1* mutant alleles, which are expected to result in loss-of-function and impact all *cped1* isoforms, exhibited no significant differences in the measures examined when compared to their respective wildtype controls, suggesting that *cped1* does not significantly contribute to these traits. We identified sequence differences in critical residues of the catalytic triad between the zebrafish and mouse orthologs of CPED1, suggesting that differences in key residues, as well as distinct alternative splicing, could underlie different functions of *CPED1* orthologs in the two species. Our studies fail to support a requirement of *cped1* in zebrafish bone and lean tissue, adding to evidence that variants at 7q31.31 can act independently of *CPED1* to influence BMD and fracture risk.

**Lay summary:** Bone mineral density (BMD) is a key indicator for predicting and diagnosing osteoporosis and fracture risk, and it has been estimated that up to 89% of variation in BMD is determined by genetics. Multiple human genetics studies have nominated *CPED1* as a potential gene underlying BMD and fracture risk heritability, however the function of *CPED1* remains poorly understood. In this study, we examined the role of *cped1* in bone by quantifying over 200 morphological measures of vertebral and craniofacial bone size, shape, and density in two different mutant lines of zebrafish in which *cped1* function was reduced or eliminated. We also examined lean tissue mass because co-heritability of this trait with BMD has also been hypothesized to involve *CPED1*. Surprisingly, despite the loss of *cped1* function, there were no significant differences between the mutant zebrafish and their respective controls. Our study therefore fails to support a role for *cped1* in bone and lean tissue, suggesting that hereditary influence on BMD and fracture risk can occur independently of *CPED1*.

## Introduction

Understanding genetic risk for osteoporosis is important to reduce this massive health burden. It has been estimated that up to 89% of the variation in bone mineral density (BMD), a key indicator for diagnosing osteoporosis and predicting fracture risk, is determined by genetics (1). Over the past several decades, a large number of human genetic studies have nominated Cadherin-like and PC-esterase Domain-containing 1 (*CPED1*) as a candidate target gene mediating BMD and fracture risk heritability. In particular, genome-wide association studies (GWAS) have identified human chromosome region 7q31.31, also known as the *CPED1*-*WNT16* locus, to be associated with BMD and fracture (2–9). Of the five genes at this locus (*WNT16*, *CPED1*, *ING3*, *FAM3C*, and *TSPAN12*), *WNT16* is the most studied and characterized (10). *In vivo* mouse studies have shown that *Wnt16* is necessary for bone mass and strength (5, 6), in part by suppressing osteoclastogenesis (11) and promoting osteoblastogenesis (12) in cortical bone. Moreover, functional studies in zebrafish have found that *wnt16* is an important regulator of bone formation, fracture susceptibility, and morphogenesis (13–15), further supporting the hypothesis that *WNT16* is a causal gene at this locus. Intriguingly, in the GWAS by Medina-Gomez et al., two independent signals were identified at this locus, with the second signal mapping to the intronic region of *CPED1* (5). These two signals were significant even after accounting for genetic linkage. It has been postulated that the presence of multiple independent signals at the *CPED1-WNT16* locus could reflect several causal variants that act on distinct genes (5, 10). More recently, Chesi et al. performed high-resolution Capture C combined with ATAC-seq in human mesenchymal stem cells (MSCs) undergoing osteoblastic differentiation and found significant interactions between the *CPED1* promoter and SNPs linked to variants associated with BMD (16). Thus, there is evidence that *CPED1* could function as a target gene at the *CPED1-WNT16* locus, possibly in tandem with *WNT16* (10).

The biological function of the encoded protein for *CPED1* is poorly characterized. Previously known as *C7orf58*, *CPED1* was identified in a search for homologs of *Cas1p*, which encodes a fungal protein that has a novel N-terminal globular domain predicted to interact with and modify glycoproteins at the cell surface (17). Named the PC-esterase domain, proteins possessing this feature are found exclusively in the eukaryotic kingdom (18). Metazoan members of the PC-esterase family were found to have three additional predicted domains fused to the N-terminus of the PC-esterase domain: a signaling peptide sequence to target the protein to the secretory pathway at the plasma membrane, a domain related to the tubulin-tyrosine ligase family, and a cadherin-like domain predicted to form a beta-sandwich structure that allows for extracellular interactions such as carbohydrate binding.

Several studies have sought to define the role of *CPED1* in bone. Medina-Gomez et al. first explored *CPED1*’s role in bone by assessing the relationship between *CPED1* expression and BMD. These authors found that high *CPED1* expression is associated with lower BMD, as evidenced by a strong inverse correlation between *CPED1* transcript levels in human iliac crest samples and total body BMD and skull BMD (5). Subsequently, Maynard et al. performed the first in-depth experimental characterization of *Cped1* transcription and identified *Cped1* transcripts in multiple mouse tissues including bone (1). Interestingly, in addition to identifying full-length transcripts, they observed multiple alternatively spliced isoforms with missing exons, including in-frame exon deletions, as well as a severely truncated transcript predicted to encode proteins missing the cadherin-like and PC-esterase domains. They also identified several alternative transcription start sites predicted to encode for proteins with N-terminal truncations. Several functional studies examining the consequences of targeted *CPED1* knockdown have also been conducted. Chesi et al. performed siRNA-knockdown of *CPED1* in primary human MSCs subjected to osteogenic differentiation (16). These authors found that *CPED1* knockdown by targeting multiple exons did not consistently alter BMP2-induced osteoblastic differentiation, suggesting that *CPED1* may not be required for osteoblastic differentiation in human MSCs. More recently, Conery et al. performed a CRISPRi screen targeting non-coding elements in human fetal osteoblast 1.19 cells (hFOBs) (19). Their results showed that CRISPRi targeting of a non-coding element nominated by BMD GWAS altered *CPED1* expression. However, siRNA knockdown of *CPED1* in hFOBs and in primary human MSCs failed to alter osteoblastic phenotypes. It is conceivable that non-coding elements regulating *CPED1* are active in multiple tissue and/or cell types, and that *CPED1* exerts its influence on BMD via its expression in other cell types within bone, or in non-skeletal tissues. Thus, to determine whether *CPED1* acts as a causal gene at the *CPED1-WNT16* BMD locus, *in vivo* studies establishing its role in bone mass and morphology are needed.

To better understand the role of *CPED1* in adult bone mass and morphology, we conducted a comprehensive genetic and phenotypic analysis of *cped1* in zebrafish, an emerging model for bone and mineral research (20–23). Recently, we described a zebrafish *cped1* loss of function mutant (15). As described in Watson et al., *cped1^w1003^* harbors a CRISPR/Cas9-induced indel that causes a frameshift and premature stop codon predicted to result in truncation of the Cped1 protein. However, several limitations in the study of Watson et al. prevented the unequivocal determination of the role of *cped1* in bone. First, because the primary purpose of analyzing *cped1^w1003^* in Watson et al. was to examine the role of *cped1* in lean tissue accrual, an in-depth phenotypic analysis of bone was not performed. Moreover, we observed normal levels of mutant *cped1* transcript in *cped1^w1003^* mutants (15). Thus, we could not rule out that truncated Cped1 protein products were functional. In this study, we overcome these limitations by characterizing a new *cped1* mutant allele, *cped1^sa20221^*, and performing deep skeletal phenotyping in both *cped1* mutant lines. We characterized more than 200 measures of adult vertebral, craniofacial, and lean tissue morphology. We also examined alternative splicing of zebrafish *cped1* and gene expression in various cell/tissue types. Our studies fail to support an essential role of *cped1* in adult zebrafish bone. Specifically, homozygous mutants for both of the analyzed *cped1* alleles, which are expected to result in loss-of-function and impact all *cped1* isoforms, exhibited no significant differences in the measures examined when compared to their respective wildtype controls, suggesting that *cped1* does not significantly contribute to these traits. We identified sequence differences in critical residues of the catalytic triad between the zebrafish and mouse orthologs of CPED1, suggesting that differences in key residues, as well as distinct alternative splicing, could underlie different functions of *CPED1* orthologs in the two species. Our studies support accumulating evidence that variants at 7q31.31 can act independently of *CPED1* to influence BMD and fracture risk.

## Materials and Methods

### Ethics statement

All studies were performed on an approved protocol in accordance with the University of Washington Institutional Animal Care and Use Committee (IACUC).

### Animal care

Zebrafish were housed at 28.5°C following a 14:10 hour light:dark cycle, and provided with a standard commercial diet. Research was carried out using mixed sex wildtype (AB) and two *cped1* mutant lines (*cped1^sa20221^*, *cped1^w1003^*). The *cped1^sa20221^* line was obtained from the Zebrafish International Resource Center (Eugene, OR). We used the following primers for genotyping *cped1^sa20221^*: F: 5’ – GCCCCTGTGCAAAGTGTTAAG – 3’, R: 5’ – CACAGAAAGAAAAATGGGCGGT – 3’. PCR was performed using standard conditions (35 cycles, annealing temperature of 58°C) (15), and products were run on high-resolution 3% agarose gels. Sanger sequencing was used to identify *cped1^sa20221^*mutant PCR products. *cped1^w1003^* mutants were previously described in Watson et al. (15). MicroCT scans of 90 days post fertilization (dpf) *cped1^w1003^* mutants and wildtype clutchmates generated in the study of Watson et al. were reanalyzed for this study; no new experiments in *cped1^w1003^* mutants were performed.

### RT-PCR and tissue-specific *cped1* expression

Total RNA was harvested from adult AB wildtype and homozygous mutant fish using homogenization in TRIzol (Invitrogen), adhering to the protocol specified by the manufacturer. To analyze *cped1* transcript expression in different zebrafish organs, two adult male and two adult female wildtype fish were euthanized in an ice bath and dissected using the methods described by Gupta et al (24). Organs were harvested and preserved in RNALater at -80°C until RNA isolation was performed. Total RNA was collected from vertebral bone, skeletal muscle, intestine, whole brain, skin, swim bladder, heart, eyes, and testes. cDNA synthesis was performed with the Superscript IV First Strand Synthesis System (ThermoFisher) and 1 uL of each cDNA was used for PCR amplification. All products were electrophoresed on a single 3% agarose gel, and bands of interest were excised from the gel for DNA extraction followed by Sanger sequencing to validate the yielded product. To test for the effects of the *cped1^sa20221^* mutation on transcript expression and splicing, we used the following primers targeting exons 14 and 16 to flank the SNP mutation in exon 15: F: 5’–GCAGAATCAACATTCTGGTGATGGA–3’, R: 5’– CAGGACTGCAGCTGAGGTTGAT–3’.

### Single-cell RNA-seq analysis

We obtained the zebrafish embryonic single-cell RNA-seq (scRNA-seq) dataset by Saunders et al. (25) from the NCBI Gene Expression Omnibus repository (GSE202639). All analyses were performed using Monocle (v3) and Seurat (v4) in R (26–28). Average expression values were determined using the AverageExpression() function, which exponentiates log normalized count data prior to calculating the average value.

### microCT scanning and analysis

Fish were scanned at 90 dpf using a vivaCT40 microCT scanner (Scanco Medical, Switzerland) with the following parameters: 21 μm voxel resolution, 55kVp, 145mA, 1024 samples, 500proj/180°, 200 ms integration time (29, 30). . Four fish were scanned simultaneously in each acquisition. Scanco software was used to generate DICOM files for each fish. Vertebral bone analysis was performed using FishCuT as outlined in (30, 31). Some measures were performed on maximum intensity projections of the DICOM images as described in (15).

To calculate lean tissue volume, we followed the procedures described in Watson et al (15). Briefly, DICOM files were opened in Fiji (32) and a single slice from the midpoint of the stack, adjacent to the posterior swim bladder, was chosen for analysis. Lower and upper thresholds were automatically determined using the Default and MaxEntropy threshold algorithms within Fiji. Subsequently, a custom MATLAB script was used to open DICOM files and perform voxel segmentation based on intensity into three tissue compartments: presumptive adipose (below lower threshold), lean (between lower and upper threshold), and bone (above upper threshold).

### Craniofacial analysis

To examine craniofacial shape variation, we employed landmarking methods we have previously described (33, 34). Thirteen landmarks (Table S1), derived from Diamond et al. 2022 (33), were manually placed on anatomical prominences of the zebrafish skull using the markups module in 3DSlicer (www.slicer.org) (35). The distance between specific pairs of points was calculated, providing a total of seven measurements: skull length (from the most anterior point of the frontal bone to the most dorsal part of the first vertebrae), frontal bone length, epiphyseal bar length, distance between anguloarticulars, dorsal skull width (distance between pterotic cartilage bones), midline skull width (distance between the most posterior part of opercles), and ventral skull width (distance between the sutures connecting the opercle and interopercle). We chose these measurements based on past studies showing morphological alterations of the opercular region and frontal and parietal bones in response to mutations in genes associated with human skeletal conditions (33, 34). For analysis, we normalized the measurements for each fish based on its standard length and performed a t-test for each measurement using the statistical analysis software, Prism (v8, GraphPad). We excluded fish exhibiting gross deformations to the skull arising from sample processing for microCT scanning, leaving 6 homozygous *cped1^w1003^*mutants, 7 homozygous *cped1^sa20221^* mutants, as well as 7 of each of their respective wildtype clutchmates.

### Multiple sequence alignment

All amino acid sequences for the multiple sequence alignment were obtained from the NCBI protein database, using the isoforms with the longest annotated sequences. The sequences were aligned using the MUSCLE alignment software (version 3.8.425) on the EMBL-EBI website. We used the ggmsa package (v1.0.3) in R to determine the consensus sequence and to plot the alignment (36).

### Statistical analysis

For most results, data are reported from a single experiment. Each biological replicate represents one technical replicate. Empirical data are shown as either individual measurements or are reported as mean ± SEM. Group sizes (n) are reported in the figure panels themselves or in respective legends. Outliers were not identified; all data were included in statistical analyses. Multivariate analysis of vertebral data using the global test was performed using the globaltest package in R (30). All other statistical analyses were performed in GraphPad Prism as described in the text. p<0.05 was considered statistically significant in all cases.

## Results

### Alternative splicing of *CPED1* orthologs is species-dependent

The zebrafish *cped1* gene is located on chromosome 4 and exhibits synteny with the human *CPED1-WNT16* locus, with *cped1* flanked on the 5’ end by *ing3* and on the 3’ end by *wnt16*. Two alternatively spliced transcripts are annotated in the zebrafish GRCz11 (GCA_000002035.4) genome assembly on ENSEMBL: *cped1*-201 and *cped1*-202 (ENSDART00000067251.6 and ENSDART00000143690.4, respectively) (Fig 1A, top). Both transcripts contain 21 exons, with exons 1, 11, 12, 13, 17, 18, 19, and 21 exhibiting differences at the 5’ or 3’ end of the exon likely due to alternative donor and acceptor sites. Notably, whereas transcript isoforms exhibiting exon skipping are annotated for mouse *Cped1* (1), no such isoforms were annotated for zebrafish *cped1*.

**Fig 1.**
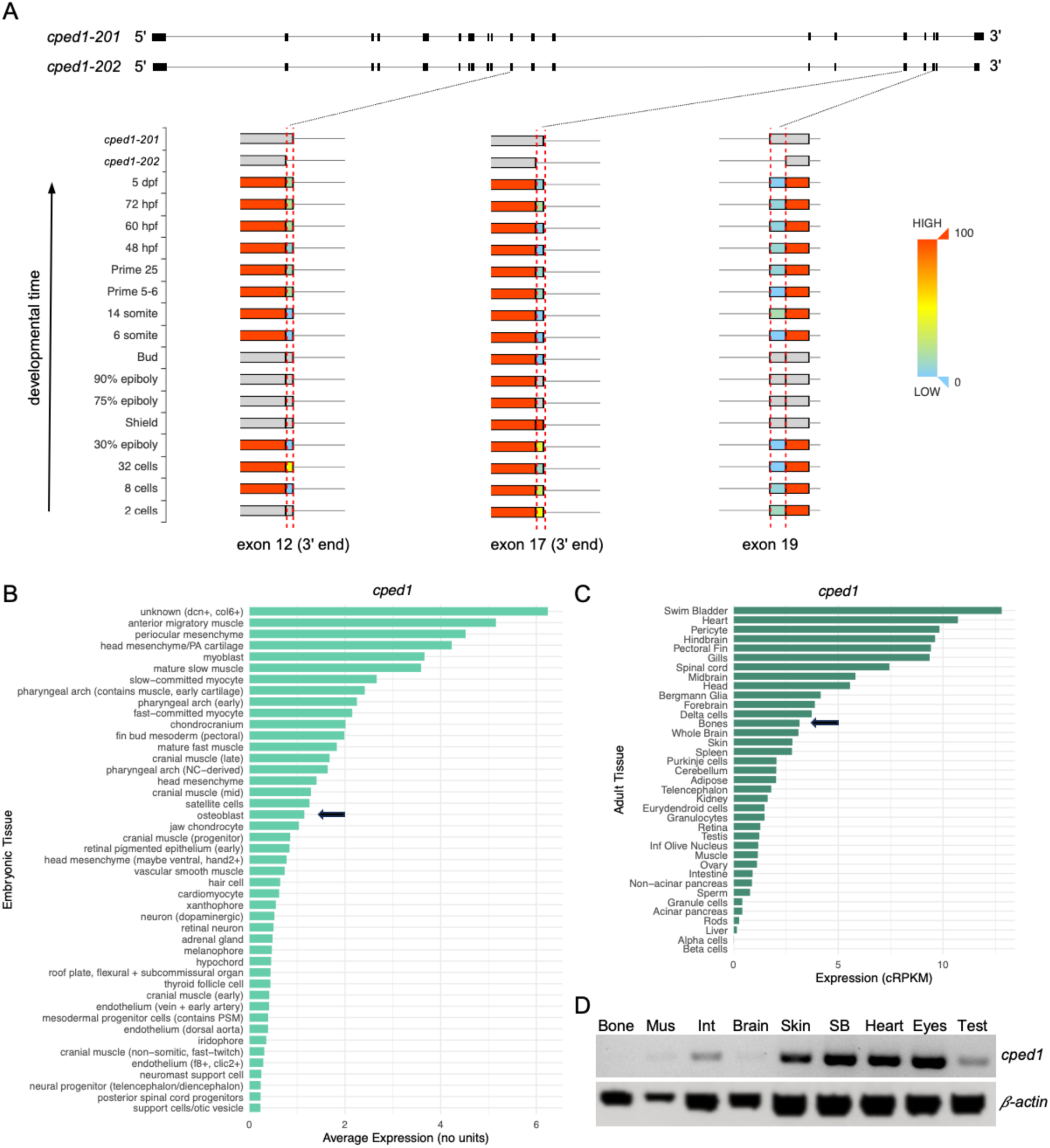
Characterization of *cped1* expression. (A) Alternative splicing events for *cped1*. Top: Exon-intron maps for the two *cped1* transcript isoforms. Bottom: Closeup views of exons 12, 17 and 19 show regions of these exons that vary in usage (denoted by the dotted red lines). The top two rows show how these exon regions differ between the two *cped1* transcript isoforms. The remaining rows show these exon regions at various stages of zebrafish development, from 2 cell stage up to 5 dpf, and are colored by usage, e.g. red indicates 100% usage in the exon, blue indicates 0% usage in the exon. Gray indicates no transcript evidence was detected at the specified developmental stage. Images for the three exonic regions were obtained from the MeDAS database (37), cropped, and assembled into a single image. (B-C) Barplots showing average *cped1* expression in multiple cell/tissue types. (B) *cped1* expression in zebrafish embryos/larvae, from the zebrafish scRNA-seq dataset published in (25). The top 45 cell populations with the highest *cped1* expression are shown. Osteoblasts are highlighted by an arrow. (C) *cped1* expression in adult zebrafish, as reported in the VASTDB database (38). Bone is highlighted by an arrow. (D) RT-PCR for *cped1* expression in tissues from wildtype adult fish. Primers amplified a region spanning exons 14 through 16. Amplification of *β-actin2* was performed in parallel and served as a loading control. Bone: Vertebral bone, Mus: Muscle, Int: Intestine, Brain: Whole brain, SB: Swim bladder, Test: Testes.

To examine alternative splice events in zebrafish *cped1*, we analyzed several public databases. First, to characterize such events in zebrafish embryos and larvae, we queried the MeDAS database, which utilizes public RNA-seq data to characterize alternative splicing events in developmental time course experiments. MeDAS quantifies inclusion of exons as well as exonic regions that contribute to different 5’ or 3’ boundaries (37). Our query identified three alternative splicing events in zebrafish *cped1* that were detected by the MeDAS pipeline, consisting of alternative exon boundary regions at the 3’ end of exon 12, the 3’ end of exon 17, and the 5’ end of exon 19 (Fig 1A, bottom). None of these alternative splice events showed significant differences in expression across developmental stages (Kruskal-Wallis p>0.05 for all three events, as determined by the MeDAS pipeline). Next, to identify alternative splice events in adult zebrafish tissues including bone, we analyzed VastDB, a large database of alternative splice events estimated using transcriptomic profiles from different cell/tissue types (38). Alternative splice events characterized by VastDB include cassette exons and microexons, alternative 5′ and 3′ splice site choices, and intron retention. Our query of VastDB revealed no alternative splice events for zebrafish *cped1* meeting a minimum threshold of alternative usage. Maynard et al. previously showed that *Cped1* exon expression varies during osteoblast differentiation, suggesting that alternative splicing differs during this process (1). Previously, our lab conducted an analysis of alternatively spliced transcripts in *sp7*+ osteoblasts during zebrafish fin regeneration (39). Using VAST-TOOLS (38) and a public RNA-seq dataset (40), we identified 1144 alternative splicing events differentially observed between 0 and 4 days post amputation (39). We searched the results of this analysis and found no alternative splicing events for *cped1*. Taken together, our analyses fail to support alternative splice events involving exon skipping for zebrafish *cped1* that are similar to events that occur for mouse *Cped1*. Thus, when referring to *cped1* gene expression below, we assume results are for transcripts harboring all 21 exons unless otherwise noted.

### *cped1* is expressed in multiple cell/tissue types

To determine *cped1* expression patterns during zebrafish embryonic development, we analyzed a publicly available single-cell RNA-sequencing (scRNA-seq) dataset by Saunders et al. (25) which profiled 1,223 zebrafish embryos collected from 19 timepoints between 18 and 96 hours post fertilization (hpf), to assess *cped1* gene expression in the 156 annotated cell clusters (Fig 1B). An unknown cluster of cells marked by *dcn*^+^/*col6*^+^ expression exhibited the highest level of *cped1* expression (average expression = 6.24). Moderate *cped1* expression in the osteoblast cell cluster was observed (average expression = 1.149). With regard to other annotated cell clusters, we found that *cped1* expression was prominent in muscle and muscle progenitor cell clusters, mesenchymal cells, and cells in the pharyngeal arch. Thus, *cped1* is broadly expressed in multiple cell types, including moderate expression in osteoblasts.

To determine *cped1* expression in adult tissues, we queried the VASTDB database (38), which collates bulk RNA-sequencing for a variety of adult zebrafish tissues. VASTDB reported the highest expression of *cped1* in swim bladder (12.80 cRPKM) and heart (10.71 cRPKM), with moderate expression of *cped1* in bone (3.15 cRPKM) (Fig 1C). Next, we examined the distribution of *cped1* expression in wildtype adult zebrafish using RT-PCR. Primers were designed to amplify a region spanning exons 14 through 16, and RT-PCR was performed on cDNA from nine solid organ tissues (Fig 1D). All PCR products were confirmed by Sanger sequencing. Highest *cped1* expression was detected in tissues from the swim bladder, heart, and eyes. We also observed moderate *cped1* expression in the skin, intestines, and testes. *cped1* expression was only faintly detectable in muscle and brain tissues, and virtually undetectable in vertebral bone. Notably, moderate levels of *cped1* were detected in bone by RNA-sequencing in VASTDB, thus it is possible that *cped1* transcript levels in vertebral bone, while present, are too low to be detected in our RT-PCR assay. Taken together, our results suggest that *cped1* is broadly expressed in multiple adult tissues in zebrafish, with moderate or low expression in bone.

### Characterization of the *cped1^sa20221^* allele

To examine whether *cped1* is necessary for bone mass and morphology, we analyzed *cped1^sa20221^* mutant zebrafish. The *cped1^sa20221^* allele was generated as part of the Wellcome Sanger Institute’s ENU mutagenesis project (41). The allele harbors a single point mutation in exon 15 that results in a premature stop codon (ENSDART00000143690.1:c.1922C>A; p.Ser641*). Exon 15 maps to a region upstream of the predicted PC-esterase domain of the protein, thus, translation of the predicted truncated transcript should result in loss of the PC-esterase domain (Fig 2A). To determine the effects of the premature stop codon on the transcript, we performed RT-PCR spanning the exons flanking the mutation-harboring exon using tissues from adult homozygous mutants (which we henceforth refer to as *cped1^sa20221^* mutants) and wildtype clutchmates. We found that the expression level of the *cped1* transcript was visibly reduced in *cped1^sa20221^*mutants compared to wildtype (Fig 2C), suggesting that the mutant transcript is degraded by nonsense-mediated decay (NMD). While CRISPR-induced indels have been shown to generate novel splice variants that can help protect against deleterious mutations (42), we did not observe evidence of exon skipping resulting in loss of exon 15 in *cped1^sa20221^* mutants. Based on the reduced *cped1* transcript in *cped1^sa20221^* mutants, as well as the predicted loss of the PC-esterase domain, we conclude that *cped1^sa20221^* is likely to be functioning as a strong hypomorph or null allele.

**Fig 2.**
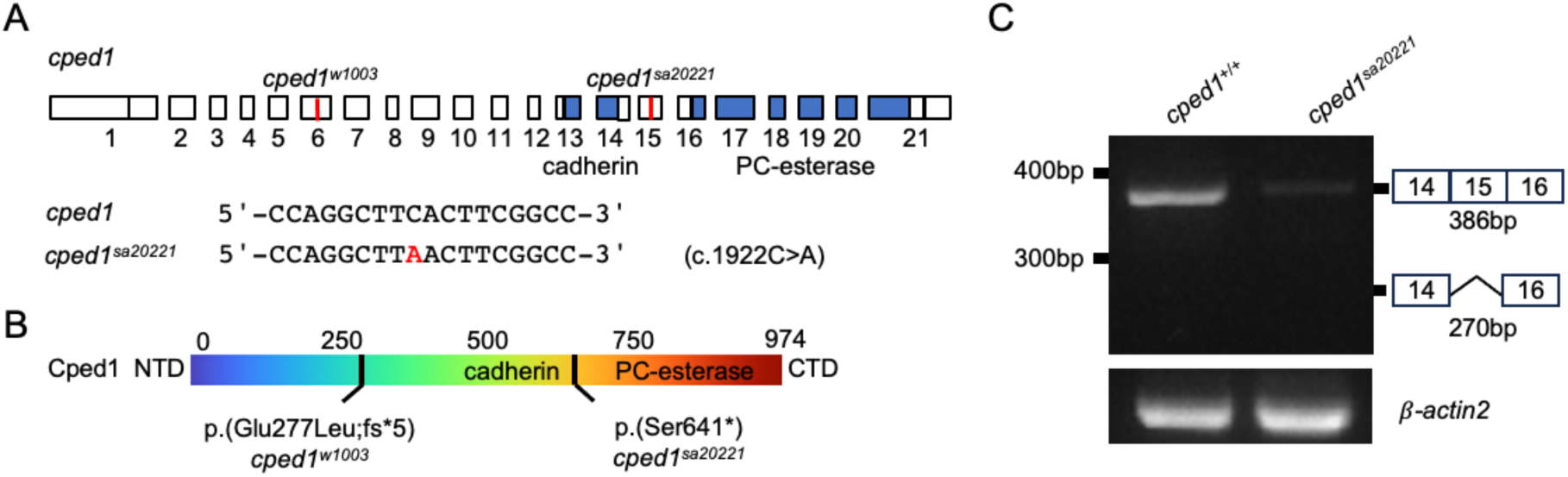
Characterization of the *cped1^sa20221^* allele. (A) Sequence and genomic location of *sa20221*. Blue highlighted regions indicate the locations of the exonic regions encoding for the cadherin-like domain and PC-esterase domain. (B) Predicted effects of *sa20221* on Cped1 amino acid sequence. For (A) and (B), information for *w1003*, an allele we previously described (15), is shown for reference. (C) Mutant mRNA is degraded in *cped1^sa20221^* mutants. RT-PCR for the region spanning exons 14 through 16 of *cped1* in adult whole body tissues shows reduced transcript levels in *cped1^sa20221^* mutants compared to controls. Evidence of exon skipping in exon 15 was not observed.

### *cped1^sa20221^* mutants exhibit normal vertebral bone mass and morphology

To determine whether *cped1* is necessary for bone mass and morphology, we performed deep vertebral bone phenotyping in adult *cped1^sa20221^*mutants. Adult *cped1^sa20221^* mutants and wildtype clutchmates were collected at 90 dpf and scanned by microCT (n=8 per group). We used FishCuT to assess 200 measures of vertebral bone mass, morphology, and mineral density (30). Briefly, vertebrae of each fish were divided into three anatomical compartments (centrum, haemal arch, and neural arch) that were analyzed for three measurements (volume, tissue mineral density (TMD), and thickness), resulting in nine distinct vertebral metrics. With the addition of centrum length, we obtained a total of 10 measurements for each analyzed vertebra (20 vertebrae in total). A standard score was computed for each measurement, and the scores were color-coded and arranged into matrices that we call “skeletal barcodes” (Fig 3A). We then plotted the 10 measurements as a function of vertebral number along the axial skeleton and calculated p-values using the global test (Fig 3B-K). No significant differences in any of the measurements were found between *cped1^sa20221^*mutants and their controls. We also did not observe obvious differences in *cped1^sa20221^*mutants and controls when assessing for gross vertebral phenotypes such as vertebrae number, rib fracture calluses, neural arch non-unions, centrum fusions, or centrum compressions (Fig 3L). Thus, our studies in *cped1^sa20221^* fail to support a role of *cped1* in adult vertebral bone mass, density, and morphology.

**Fig 3.**
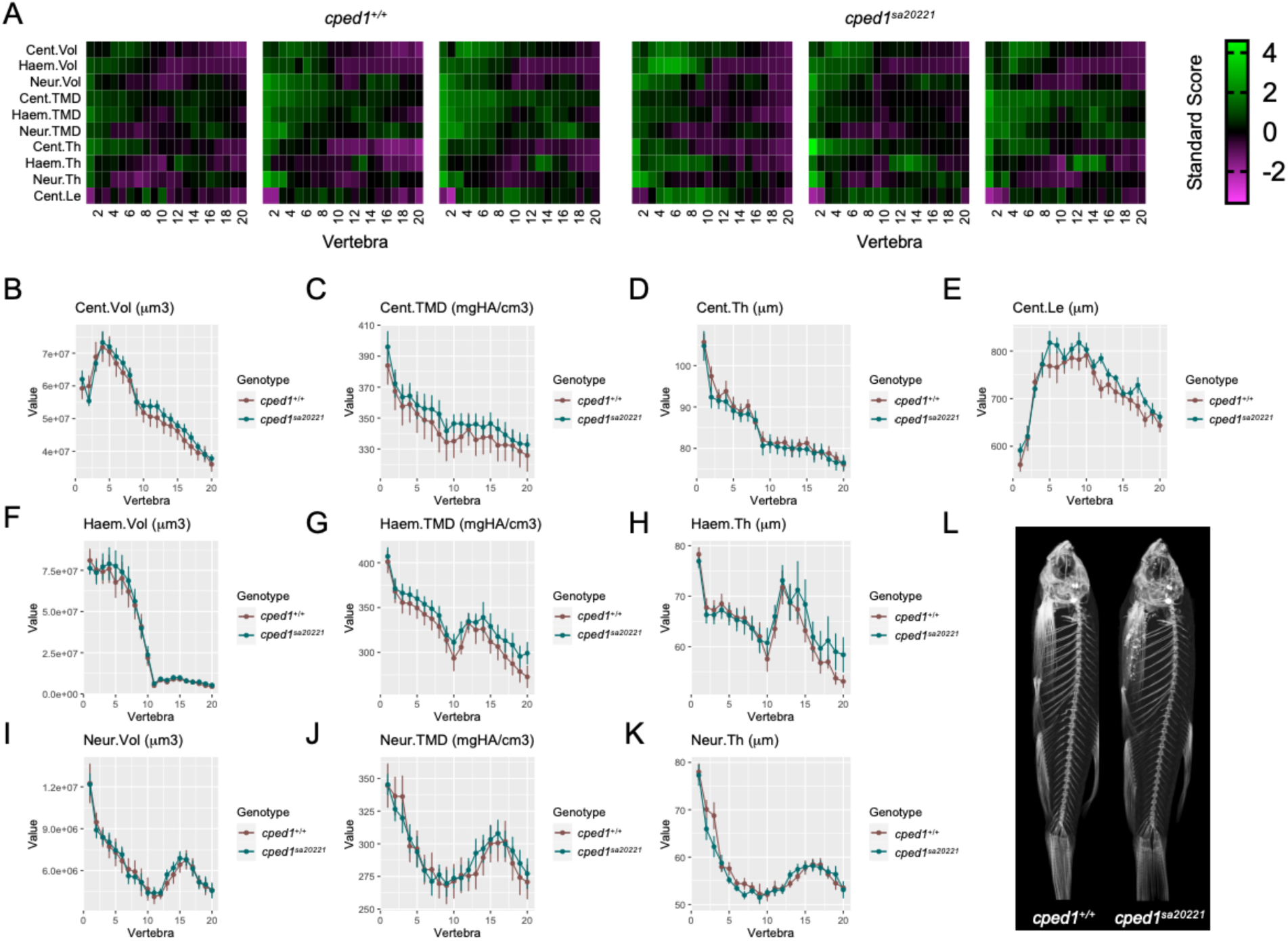
*cped1^sa20221^* mutants exhibit normal vertebral bone mass and morphology. (A) Skeletal barcodes for *cped1^+/+^* and *cped1^sa20221^* mutants (3 fish/group shown) visually depict individual vertebral phenomes. (B–K) Vertebral phenotypic measures (indicated by the graph title, with units for y axis) plotted as a function of vertebra along the spine. Values are depicted mean ± SEM (n=8/group). No measures with p<0.05 in the global test were detected. Cent, centrum; Haem, haemal arch; Neur, neural arch; Vol, volume; TMD, tissue mineral density; Th, thickness; Le, length. (L) Maximum intensity projections of microCT scans.

### Loss of *cped1* does not affect vertebral bone mass and morphology in multiple mutant lines

While our studies did not reveal a vertebral phenotype in *cped1^sa20221^*mutants, it is possible that the allele has unintended consequences that might obscure the function of Cped1. In particular, it has previously been shown that transcriptional adaptation caused by mutant mRNA degradation, a form of genetic compensation, can produce mild or even an absence of expected phenotypes (43). Thus, we sought to repeat our analyses using a different *cped1* mutant allele in which transcriptional adaptation caused by mutant mRNA degradation was unlikely. For this, we analyzed *cped1^w1003^*mutants (15). As described in Watson et al., *cped1^w1003^* harbors a CRISPR/Cas9-induced net 11 bp deletion (c.831-844delinsACT). This indel mutation causes a frameshift and premature stop codon in exon 6 (ENSDARP00000067250.4, p:Gln278Leu;fs*5) which is predicted to result in truncation of the Cped1 protein and loss of both the cadherin-like and PC-esterase domains (Fig 2B). Notably, normal levels of mutant *cped1* transcript are observed in *cped1^w1003^* mutants, suggesting the absence of transcriptional adaptation caused by mutant mRNA degradation (15). While it is possible that truncated Cped1 protein products expressed in *cped1^w1003^* mutants retain biological activity, we surmised that comparing phenotypes with *cped1^sa20221^* mutants might help to rule out this possibility. As mentioned previously, in the prior study of Watson et al., we showed that adult *cped1^w1003^* mutants do not exhibit significant differences in lean tissue-related traits when compared to their wildtype siblings (15). However, deep vertebral phenotyping using FishCuT in *cped1^w1003^* mutants was not performed (15).

We therefore conducted FishCuT vertebral analysis in adult (90 dpf) *cped1^w1003^* mutants. We observed no significant differences between c*ped1^w1003^* mutants and their wildtype clutchmates (Fig S1). Next, we combined data for c*ped1^w1003^* and *cped1^sa20221^* mutant lines in a two-way ANOVA analysis (Fig 4). We surmised that combining data for both lines into a single analysis would increase our power to detect smaller effect sizes. For this, we used the average of each of the 10 FishCuT measurements across the 16 rostral-most measured vertebrae for both mutant alleles (we chose to analyze 16 rather than 20 vertebrae for this analysis because only 16 vertebrae were analyzed for *cped1^w1003^* mutants in (15)). The two-way ANOVA allowed us to simultaneously determine whether (i) there were common phenotypic differences between wildtype and mutants for both alleles (indicated by p-value for genotype), (ii) there were baseline phenotypic differences for different alleles due to genetic background and/or environmental conditions when testing each allele (indicated by p-value for allele), and (iii) mutants for each *cped1* allele exhibited different phenotypic responses when compared to wildtype (indicated by p-value for genotype:allele interaction).

**Fig 4.**
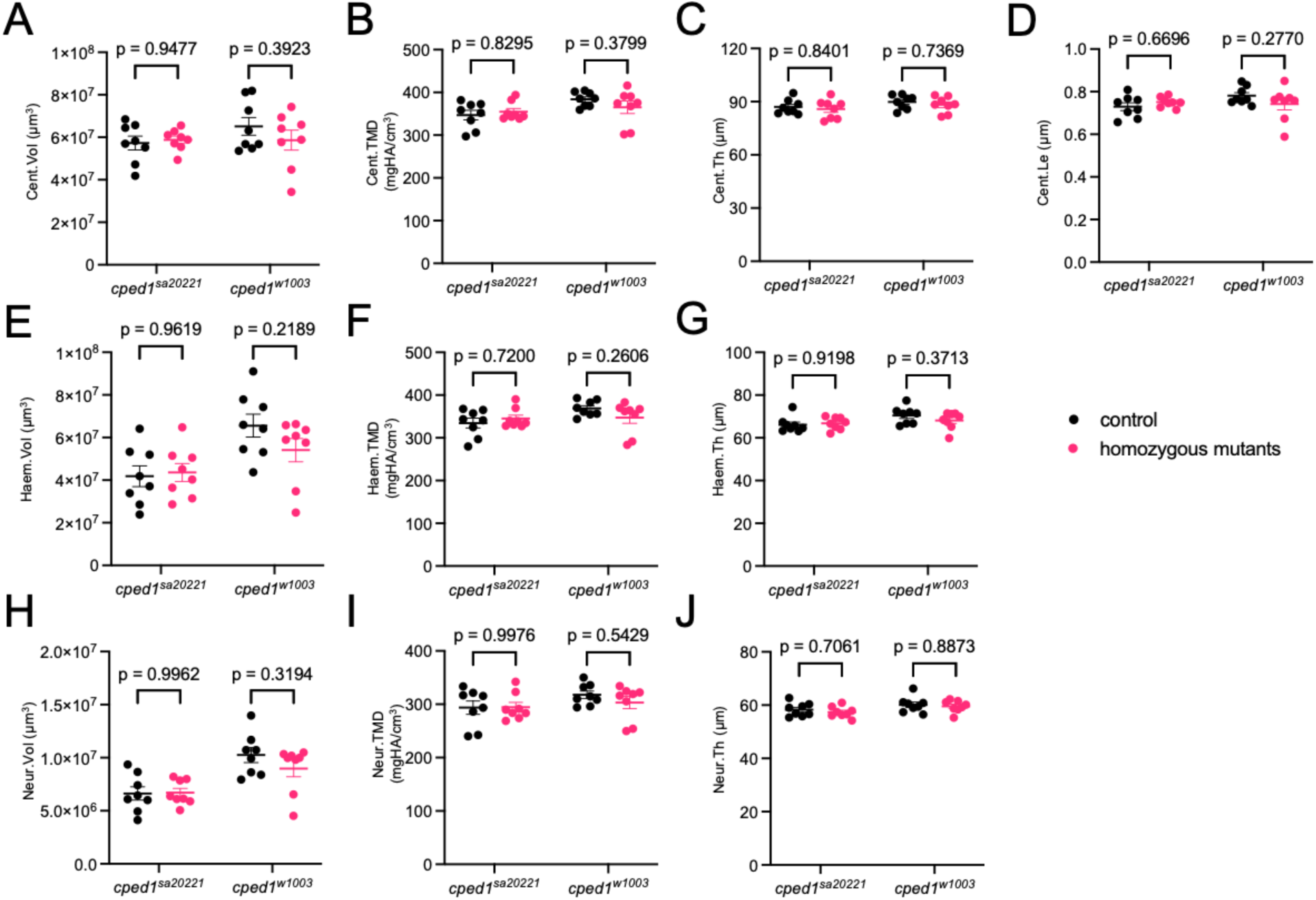
Loss of *cped1* does not affect vertebral bone mass and morphology in multiple mutant lines. (A-J) Combined analysis of vertebral measurements for *cped1^sa20221^* and *cped1^w1003^* mutants compared to their respective wildtype clutchmates. Each point represents a single fish, and the value is the average of 16 measured vertebrae. Bars indicate mean ± SEM. Shown are p-values from Sidak’s multiple comparisons tests; no significant p-values for comparisons between homozygous mutants and respective controls were observed.

The result of the two-way ANOVA indicated that there were no significant effects of genotype (i.e., wildtype versus mutant) for any of the vertebral measurements (Table 1). Moreover, Sidak’s multiple comparisons analysis for each allele found no significant differences between wildtype and mutant for either allele. We also observed no statistically significant interactions between allele and genotype, indicating that mutants for each *cped1* allele exhibited similar phenotypic responses when compared to wildtype. Several measures had p-values <0.05 for the allele factor, indicating there were baseline phenotypic differences for different alleles. Because the *cped1^w1003^* fish was generated in an AB background, whereas the *cped1^sa20221^*fish was generated in a longfin background, significant effects of allele are likely attributable to (i) background differences between the two lines and/or (ii) differences in environmental conditions that equally impacted controls and mutants for each allele (e.g., different rates of developmental progress). In sum, our studies in multiple *cped1* mutant lines fail to support a role of *cped1* in vertebral bone mass and morphology.

**Table 1.** P-values from two-way ANOVA of vertebral measures in *cped1^sa20221^* and *cped1^w1003^* mutants and respective wildtype controls.

| Measurement | Genotype | Allele | Interaction |
| --- | --- | --- | --- |
| centrum volume | p=0.503 | p=0.298 | p=0.283 |
| centrum thickness | p=0.3901 | p=0.112 | p=0.903 |
| centrum TMD | p=0.612 | p=0.029* | p=0.207 |
| centrum length | p=0.638 | p=0.262 | p=0.117 |
| haemal arch volume | p=0.3403 | p=0.0020* | p=0.1966 |
| haemal arch thickness | p=0.5184 | p=0.0339* | p=0.251 |
| haemal arch TMD | p=0.582 | p=0.0816 | p=0.1229 |
| neural arch volume | p=0.3609 | p=<0.0001* | p=0.3076 |
| neural arch thickness | p=0.4067 | p=0.027* | p=0.8259 |
| neural arch TMD | p=0.5106 | p=0.1223 | p=0.4574 |

\* = p-value < 0.05

### Loss of *cped1* does not affect lean tissue mass and morphology

Because variants at the *CPED1-WNT16* locus have been associated with pleiotropic effects on both bone and lean mass-related traits (15, 44), we also examined whether *cped1^sa20221^* mutants exhibited any phenotypic differences in lean tissue mass and morphology. For this, we measured the following traits: standard length and fineness ratio to assess body shape; total, anterior and posterior trunk lean volumes; and anterior and posterior swim bladder chamber length (15). We also manually measured centrum length, neural arch length, and neural arch angle, respectively, since these measures are highly correlated to myomere length, height, and angle, respectively (45). We previously found no significant differences in these measures in *cped1^w1003^*mutants (15). We combined data for both *cped1^sa20221^*and *cped1^w1003^* and performed a two-way ANOVA analysis. There were no statistically significant effects of genotype in any of the muscle morphology measures between controls and germline mutants (Fig 5). Furthermore, for each allele, Sidak’s multiple comparisons tests did not detect significant differences between wildtype and mutant. There were also no significant genotype:allele interactions for any of the measures. Finally, statistically significant p-values for the allele factor were detected for several measures, which we again attributed to the different backgrounds of the two lines and/or environmental differences when testing each allele (Table 2). Taken together, our findings fail to support a role of *cped1* in lean tissue mass and morphology.

**Fig 5.**
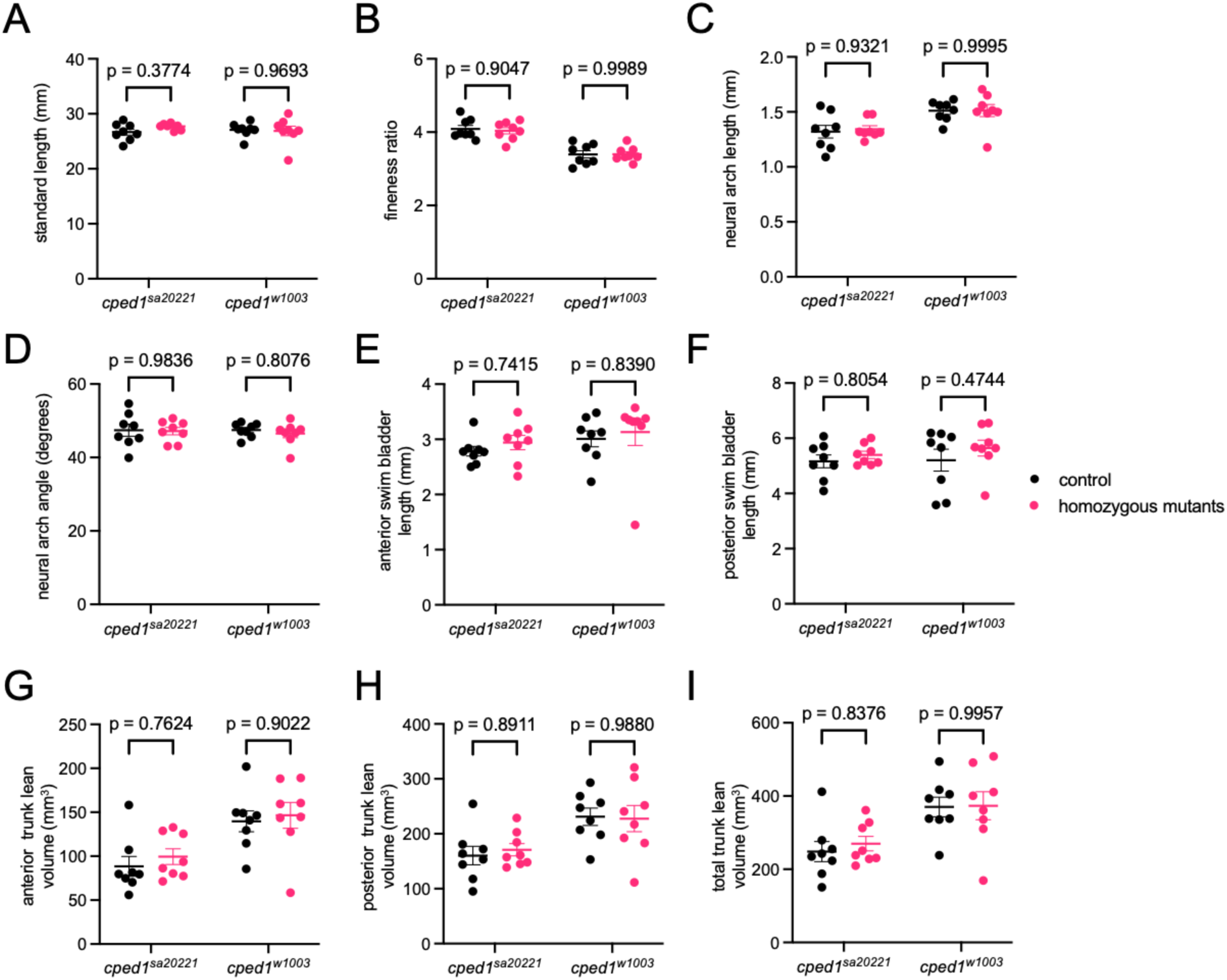
Loss of *cped1* does not affect lean mass morphology. (A-I) Combined analysis of lean mass-related measures for *cped1^sa20221^* and *cped1^w1003^*mutants and their corresponding wildtype clutchmates. Each point represents a single fish. Bars indicate mean ± SEM. Shown are p-values from Sidak’s multiple comparisons tests; no significant p-values for comparisons between homozygous mutants and their respective controls were observed.

**Table 2.** P-values from two-way ANOVA of lean mass-related traits in *cped1^sa20221^* and *cped1^w1003^* mutants and respective wildtype controls.

| Measurement | Genotype | Allele | Interaction |
| --- | --- | --- | --- |
| standard length | p=0.4611 | p=0.7151 | p=0.2968 |
| fineness ratio | p=0.8016 | p=<0.0001* | p=0.7561 |
| neural arch length | p=0.798 | p=0.0005* | p=0.8302 |
| neural arch angle | p=0.5998 | p=0.7910 | p=0.7660 |
| ant. swim bladder chamber | p=0.3921 | p=0.2103 | p=0.9082 |
| pos. swim bladder chamber | p=0.238 | p=0.6046 | p=0.7150 |
| ant. trunk lean volume | p=0.4556 | p=0.0003* | p=0.8574 |
| pos. trunk lean volume | p=0.8381 | p=0.0011* | p=0.6902 |
| total trunk lean volume | p=0.6658 | p=0.0005* | p=0.7515 |

\* = p-value < 0.05

### Loss of *cped1* does not affect craniofacial morphology

Several GWAS have suggested that variants in the intronic region of *CPED1* may influence human craniofacial morphology. For instance, Medina-Gomez et al. found that variants most strongly associated with skull BMD were located in the intronic region of *CPED1*, with rs7801723 identified as the most significantly associated SNP (5). Moreover, in a GWA study identifying human genomic loci associated with craniofacial morphology in a Latin American population, Bonfante et al. detected a single nucleotide variant in the intronic region of the *CPED1* gene (rs6950680) associated with “jaw protrusion” (46). To assess whether *cped1* zebrafish mutants exhibited altered craniofacial morphology, we performed landmark-based morphometric analyses using methods we have previously described (33, 34). Using the Markups module in 3D Slicer (35), we manually placed 13 markers on distinctive craniofacial landmarks of microCT scans of *cped1^sa20221^* and *cped1^w1003^* mutants and their respective wildtype clutchmates, and measured the distances between 7 pairs of landmarks (Fig. 6A, B). From our two-way ANOVA analysis, we did not find any significant effects of genotype for any of the craniofacial measurements (Fig. 6C-I and Table 3). In addition, Sidak’s multiple comparisons test did not detect significant differences between homozygous mutants and their respective wildtype controls for either allele. Moreover, no significant interactions were found between genotype and allele. For reasons described previously, statistically significant p-values for the allele factor were detected for several measures similar to previous analyses. In sum, our findings fail to support a role of *cped1* in craniofacial morphology.

**Fig 6.**
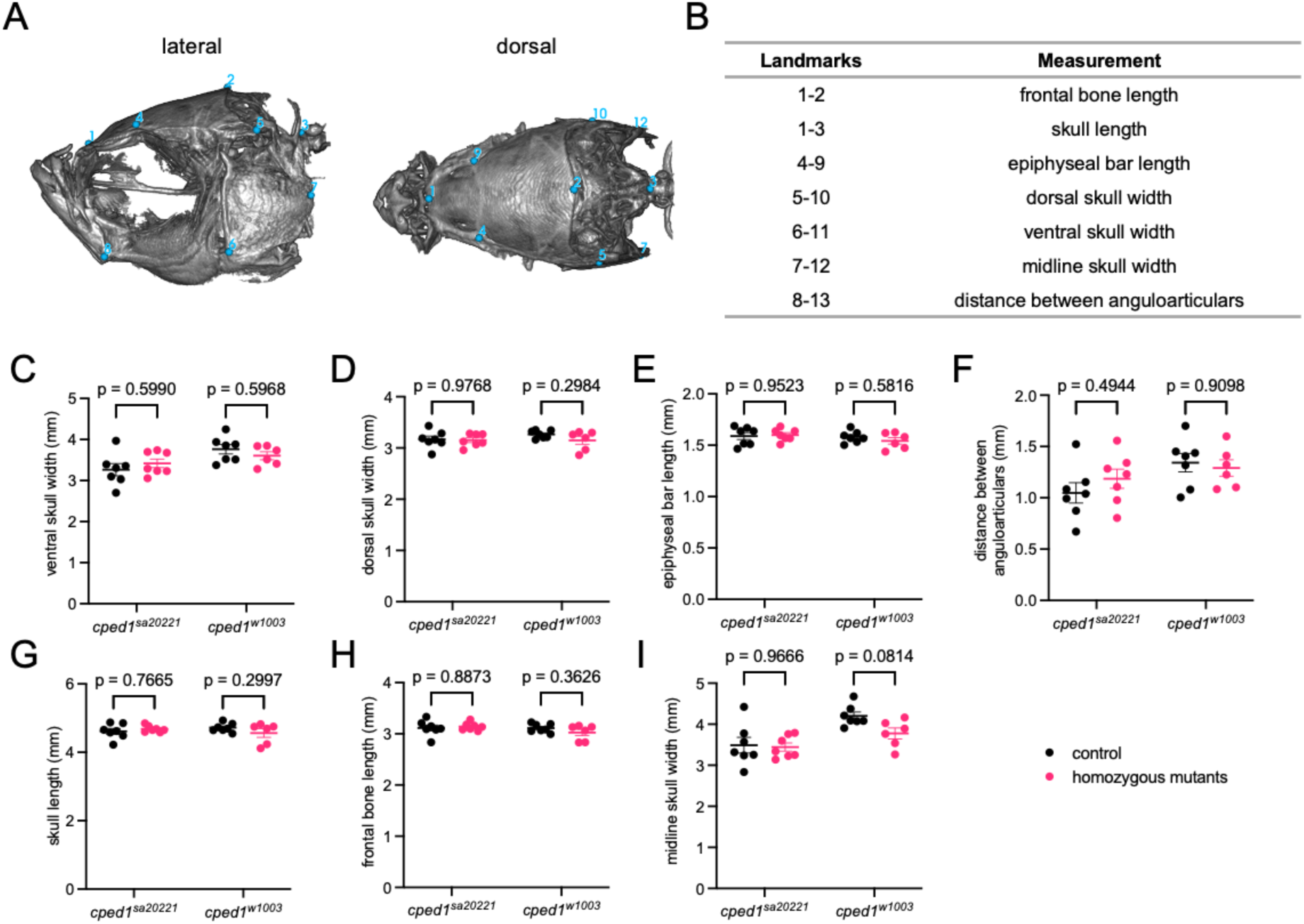
Loss of *cped1* does not affect craniofacial morphology. (A) Anatomical landmarks used for measurements. Shown are the lateral view (left) and dorsal view (right) of a wildtype zebrafish skull. Landmark positions are labeled in blue. Note that landmark 11 is on the right lateral surface, opposite landmark 6, and landmark 13 is on the right lateral surface, opposite landmark 8. (B) Distances between pairs of landmarks were used as measurements to quantify craniofacial morphology. (C-I) Combined analysis of craniofacial morphological measures for *cped1^sa20221^* and *cped1^w1003^*mutants and their corresponding wildtype clutchmates. Each point represents an individual fish. Shown are p-values from Sidak’s multiple comparisons tests; no significant p-values for comparisons between homozygous mutants and controls were observed.

**Table 3.** P-values from two-way ANOVA of craniofacial phenotyping comparison of *cped1^sa20221^* and *cped1^w1003^* and their respective wildtype controls.

| Measurement | Genotype | Allele | Interaction |
| --- | --- | --- | --- |
| skull length | p=0.566 | p=0.922 | p=0.1485 |
| frontal/parietal bone length | p=0.526 | p=0.218 | p=0.223 |
| epiphyseal bar length | p=0.629 | p=0.243 | p=0.389 |
| distance between anguloarticulars | p=0.643 | p=0.0409* | p=0.312 |
| midline skull width | p=0.099 | p=0.0009* | p=0.177 |
| dorsal skull width | p=0.252 | p=0.404 | p=0.374 |
| ventral skull width | p=0.976 | p=0.0089* | p=0.205 |

\* = p-value < 0.05

### The PC-esterase domain in zebrafish Cped1 lacks a conserved catalytic domain

Our results so far showed negligible effects on bone and lean mass morphology arising from two different nonsense mutations in *cped1*. This led us to ask whether the functional activity of the PC-esterase domain is conserved in zebrafish. We therefore decided to compare the Cped1 amino acid sequence of zebrafish to CPED1 orthologs in other chordates. We used the MUSCLE alignment software to perform a multiple sequence alignment of CPED1 orthologs identified from the NCBI Orthologs database: human (NP_079189.4), mouse (*Mus musculus*, NP_001074820.1), chicken (*Gallus gallus*, XP_416005.4), frog (*Xenopus tropicalis*, XP_031754845.1), lizard (*Sceloporus undulatus*, XP_042324422.1), amphioxus (*Branchiostoma floridae*, XP_035676865.1), little skate (*Leucoraja erinacea*, XP_055506810.1), and another teleost, Japanese medaka (*Oryzias latipes*, XP_023808151.1). We also performed a BLASTp search for a Cped1 ortholog in the basal chordate *Ciona intestinalis* in the non-redundant protein sequences database on NCBI but did not recover any potential orthologs. The alignment showed that, compared to the human and mouse orthologs, zebrafish Cped1 shares 39.87% and 38.92% identity over the entire protein sequence, respectively. Within the predicted PC-esterase domain, zebrafish Cped1 shares 55.6% and 54.28% identity with human and mouse orthologs, respectively (Fig S2), indicating that the PC-esterase domain is more highly conserved between zebrafish and mammals than other regions of the protein.

We then asked whether the predicted enzymatic active site of the PC-esterase domain was intact in zebrafish. To date, the crystal structure and biochemical characterization of only one member of the PC-esterase domain family has been determined, XOAT1 from *Arabidopsis thaliana* (18, 47). The crystal structure of XOAT1 revealed the presence of a canonical “catalytic triad” of amino acids adjacent to the active site (Serine-Aspartate-Histidine), which was required for the enzymatic activity of the protein as determined by site-directed mutagenesis studies. The biochemical mechanism of the catalytic triad is well characterized and typically consists of a serine (S) in a GDS motif, and an aspartate (D) and histidine (H) in a DxxH motif found upstream of the last predicted helix (17, 48). Anantharaman and Aravind previously showed that, within this catalytic triad, the serine and histidine (but not aspartate) showed absolute conservation amongst representatives of the PC-esterase family (17). Using the positions of the predicted catalytic residues, we identified the corresponding residues in our alignment (asterisks in Fig S2) and found that the predicted catalytic triad appears to be disrupted in zebrafish. Specifically, we observed that the histidine residue which showed absolute conservation amongst the PC-esterase family in the study of (17) was substituted with a glutamine (Q) in zebrafish. This substitution was not seen in any other sequence we analyzed. During ester bond cleavage, the histidine in the catalytic triad facilitates the chemical reaction by removing a proton from the serine, thereby activating the serine and allowing it to attack the ester bond (18, 49). It is therefore plausible that loss of the histidine in zebrafish disrupts the enzymatic activity of the PC-esterase domain in Cped1.

## Discussion

### Do *CPED1* orthologs have different roles in zebrafish and mouse bone?

Our analyses indicate potential differences between zebrafish and mouse *CPED1* orthologs at the gene expression and protein levels. At the level of gene expression, mouse *Cped1* exhibits exon skipping as well as expression of N- and C-terminally truncated isoforms (1). In contrast, alternative splicing of zebrafish *cped1* appears to primarily consist of alternative donor/acceptor sites. Because our analyses fail to support alternatively spliced *cped1* transcripts with missing exons, both mutant alleles in our study are expected to impact all *cped1* isoforms, allowing for an unambiguous assessment of the role of *cped1* in bone. This is in contrast to exon skipping observed for mouse *Cped1*, which makes the generation of CRISPR-induced knockout alleles more challenging. We acknowledge that, if exon skipping is critical to *CPED1*’s regulation of bone in humans, the absence of this in zebrafish *cped1* could make it an inappropriate model. In this context, it is worthwhile to note that exon skipping for exon 3 in mouse *Cped1*, which is catalogued in the VASTDB and MeDAS databases, is not predicted by VASTDB to be conserved in the orthologous human or zebrafish exons. At the level of the encoded protein, we found that zebrafish Cped1 is predicted to harbor an amino acid substitution for a key residue within the catalytic triad of the PC-esterase domain, a substitution which is not observed in mouse, human, or any other representative of the PC-esterase family in the study of (17). Even though the functional domains of CPED1 have not yet been determined experimentally, the loss of a putative catalytic amino acid in the predicted PC-esterase domain of zebrafish Cped1 suggests that the protein’s function may have been lost during evolution. Taken together, there are predicted transcriptional and protein differences in *CPED1* zebrafish and mouse orthologs that could underlie their different roles in bone.

### Does *CPED1* influence BMD and fracture risk heritability at the *CPED1-WNT16* locus?

While we acknowledge that the role of zebrafish *cped1* in bone may be different than the role of *CPED1* in mammals, it is worthwhile to note that our findings share some similarities with previous functional studies. In particular, our studies fail to support an essential role of *cped1* in bone, which is in agreement with studies by Chesi et al. who found that *CPED1* knockdown did not alter BMP2-induced osteoblastic differentiation in primary human MSCs (16). Our studies also support the studies of Conery et al. who found that *CPED1* knockdown did not alter osteoblastic phenotypes in primary human MSCs and hFOBs (19). In addition, *Cped1* mutant mice analyzed by the International Mouse Phenotyping Consortium did not exhibit any significant bone phenotypes (www.mousephenotype.org) (50). These mice also did not exhibit any significant differences in lean mass or craniofacial-related measures. These studies bring forth the question of whether the *CPED1-WNT16* locus could influence BMD and fracture independently of *CPED1*. Other genes at 7q31.31 (*ING3*, *FAM3C*, and *WNT16*) have previously been shown to have a role in regulating bone morphology and strength and/or osteoblast differentiation. For example, Chesi et al. found that siRNA-mediated knockdown of *ING3* in primary hMSCs reduced BMP2-induced osteogenic differentiation, suggesting that ING3 is needed for osteoblast differentiation (16). In studies by Maata et al. and Bendre et al., *FAM3C* was shown to be important for bone morphology in mouse, and in mediating alkaline phosphatase expression during osteoblastic differentiation (51, 52). Finally, a large body of evidence has shown that *WNT16* is likely a causal gene at the locus. For example, *Wnt16* is required for cortical bone mass and strength in mice (5, 6, 11, 12, 53–57), and in humans, expression levels of *WNT16* from iliac crest samples were positively correlated with measures of BMD at multiple skeletal sites including total body, skull, legs, total hip, and lumbar spine (5). In addition, higher levels of *WNT16* expression were correlated with higher total body lean mass (TBLM) and BMD, suggesting that *WNT16* has pleiotropic effects on BMD and lean mass. This finding was supported by a bivariate GWAS meta-analysis in a pediatric cohort that identified the *CPED1-WNT16* locus as harboring the lead variant most significantly associated with TBLM and total body less head-BMD. This finding was also supported by our recent study showing that (i) *wnt16* is specifically expressed in dermomyotome and notochord, structures critical for the development of muscle and the spine, respectively, and (ii) *wnt16*^-/-^ zebrafish mutants exhibited altered vertebral bone morphology and lean mass and morphology (15). Thus, although it cannot be ruled out that *CPED1* influences bone and lean mass in humans, there are multiple candidate genes in this locus that could play an equal or larger role in influencing musculoskeletal traits.

### Limitations

There are some limitations to our study. First, while we found no effects of loss of *cped1* in 90 dpf adult bone, it cannot be ruled out that its loss affects bone at a later stage (e.g., in aged animals). It is also possible that *cped1* plays a critical role in tissues and structures in zebrafish that we did not analyze. Additional analyses of other tissues are warranted, especially in tissues exhibiting strong *cped1* expression such as the heart and swim bladder. Second, as mentioned previously, it is possible that genetic compensation could reduce the effects of *cped1* loss in zebrafish. Our studies do not support genetic compensation arising from transcriptional adaptation due to mutant mRNA degradation or removal of the PTC-harboring exon (exon skipping). However, other forms of genetic compensation can occur, for example, through cryptic splice sites, cryptic start sites, or intron inclusion (58–64). While we cannot exclude these possibilities, our analysis of two independent *cped1* zebrafish lines harboring mutations in different exons makes it less likely that our results are due to the same compensatory effects. Lastly, our studies were not designed to include sex as a biological variable, precluding us from determining whether results differed with sex. However, it is noteworthy that the association between BMD and the *CPED1-WNT16* locus does not show significant evidence of sex-specificity (4), suggesting that if *CPED1* acts as a causal gene at the locus, the mechanism by which it contributes to BMD is unlikely to exhibit significant sexual dimorphism.

## Conclusion

In conclusion, our studies fail to support a role of *cped1* in contributing to adult zebrafish bone mass, lean mass, or bone and lean tissue morphology. Because of differences in key residues as well as distinct alternative splicing in zebrafish and mouse orthologs of *CPED1*, in-depth bone phenotypic analyses in *Cped1* knockout mice are warranted. Moreover, there is evidence that variants at 7q31.31 can act independently of *CPED1* to influence BMD and fracture risk, including recent *in vitro* studies showing that loss of *ING3* disrupts osteoblast differentiation in primary human MSCs (16). Thus, further *in vivo* studies examining the role of other genes at the locus are needed.

## Acknowledgements

Research reported in this publication was supported by the National Institute of Arthritis and Musculoskeletal and Skin Diseases of the National Institutes of Health under Award Number AR074417 as well as the Office of the Director (OD). AEG was supported by the National Institute on Aging of the National Institutes of Health, under Award Number AG066574, the Biological Mechanisms for Healthy Aging Training Grant. The content is solely the responsibility of the authors and does not necessarily represent the official views of the National Institutes of Health. The authors have no relevant financial or non-financial interests to disclose.

## Data Availability Statement

The data are available from the corresponding author upon reasonable request.

