## Supplemental Materials for "Loss of *cped1* does not affect bone and lean tissue in zebrafish"

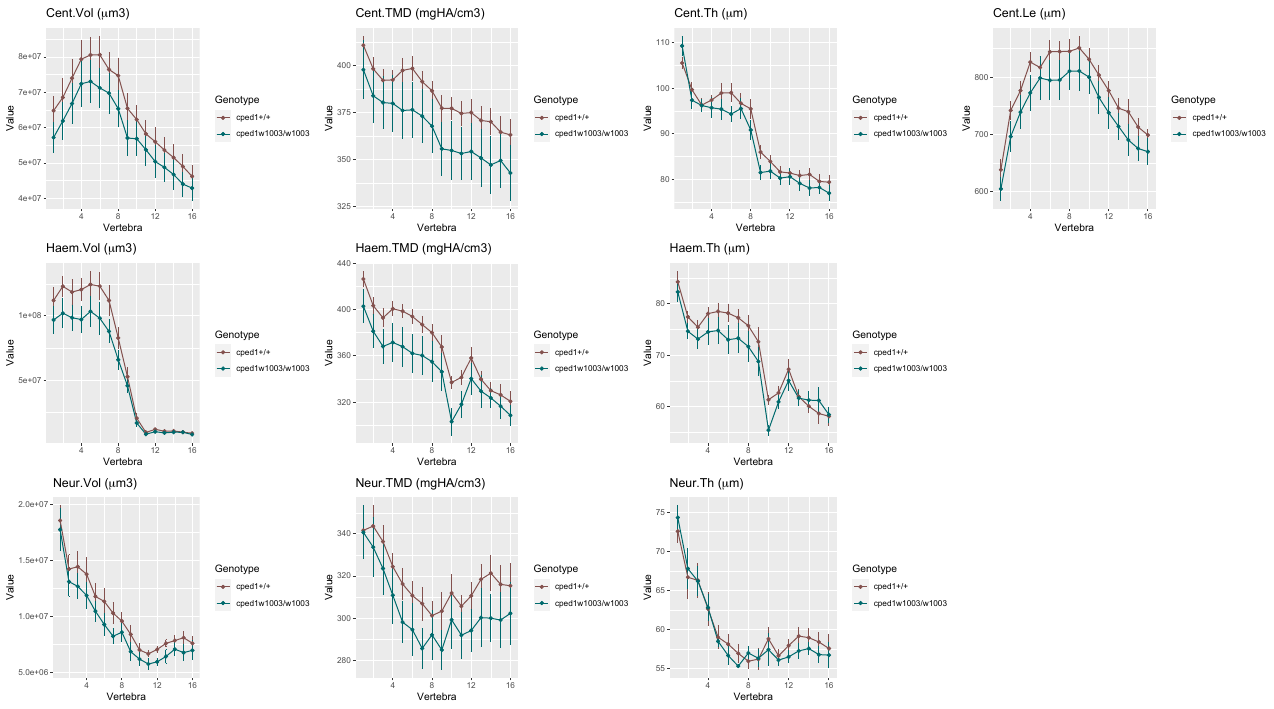

**Fig S1. *cped1^w1003^* mutants exhibit normal vertebral bone mass and morphology*.*** Vertebral phenotypic measures (indicated by the graph title, with units for y axis in parentheses) plotted as a function of vertebra along the spine. Values are depicted as mean ± SEM (n=8/group). No measures with p<0.05 in the global test were detected. Cent, centrum; Haem, haemal arch; Neur, neural arch; Vol, volume; TMD, tissue mineral density; Th, thickness; Le, length

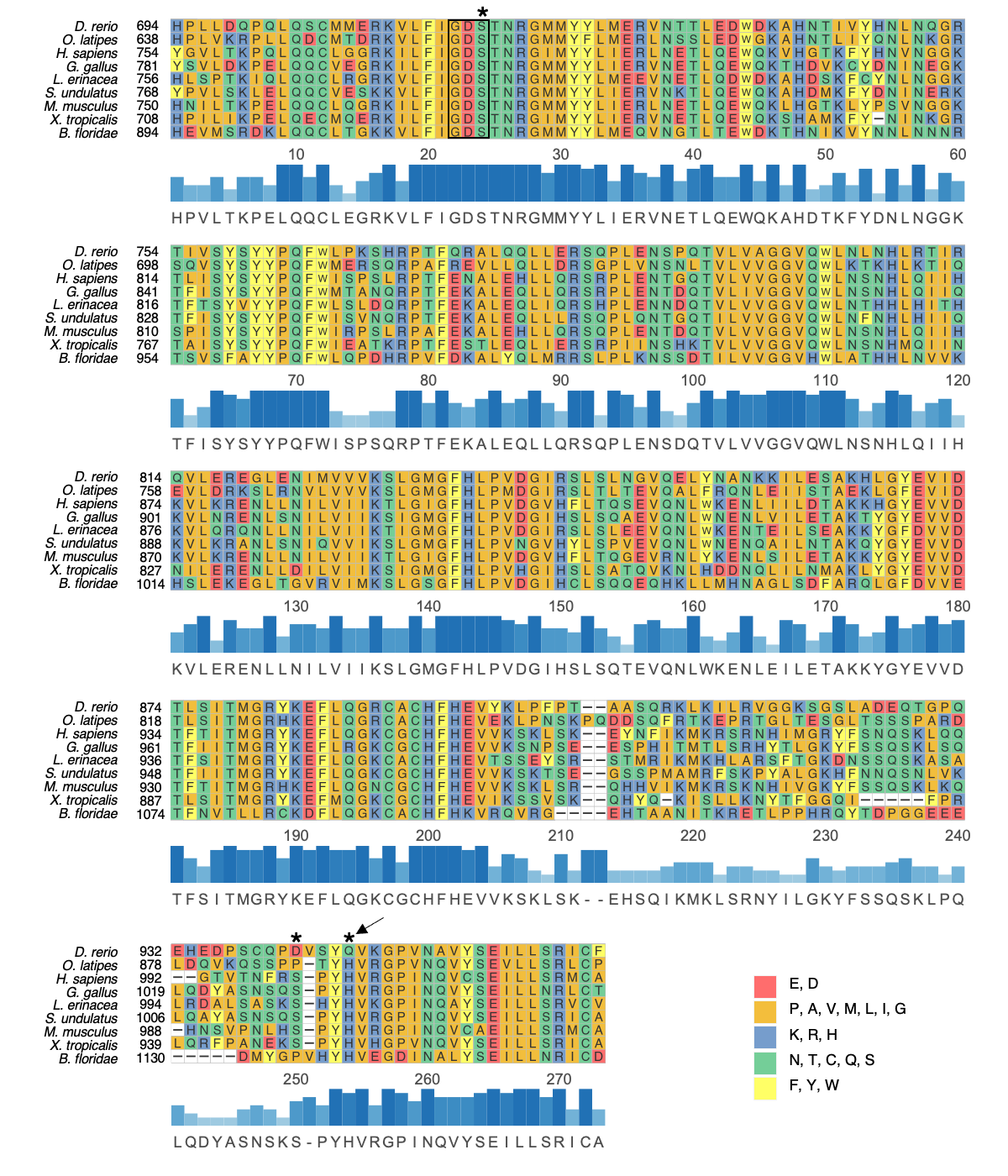

**Fig S2. Multiple sequence alignment of the predicted PC-esterase domain of CPED1 orthologs from chordate species.** Individual residues are colored based on side chain chemistry. The consensus sequence is shown below the alignment and is composed of the amino acid residues with the highest frequency at each position, with the most highly conserved residues denoted by darker, taller bars. The catalytic triad of amino acid residues is denoted with asterisks, and the GDS motif is indicated with a black box. The histidine residue which showed absolute conservation amongst the PC-esterase family in the study of [17] and which was substituted with a glutamine (Q) in zebrafish (*D. rerio*) is indicated with an arrow.

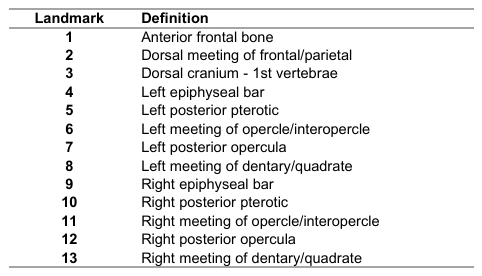

**Table S1. Anatomical descriptions of the zebrafish craniofacial landmarks used for analysis.**
